# Behaviour-driven *Arc* expression is greater in dorsal than ventral CA1 regardless of task or sex differences

**DOI:** 10.1101/2021.03.17.435850

**Authors:** J. Quinn Lee, Rebecca McHugh, Erik Morgan, Robert J. Sutherland, Robert J. McDonald

## Abstract

Evidence from genetic, behavioural, anatomical, and physiological study suggests that the hippocampus functionally differs across its longitudinal (dorsoventral or septotemporal) axis. Although, how to best characterize functional and representational differences in the hippocampus across its long axis remains unclear. While some suggest that the hippocampus can be divided into dorsal and ventral subregions that support distinct cognitive functions, others posit that these regions vary in their granularity of representation, wherein spatial-temporal resolution decreases in the ventral (temporal) direction. Importantly, the cognitive and granular hypotheses make distinct predictions on cellular recruitment dynamics under conditions when animals perform tasks with qualitatively different cognitive-behavioural demands. The cognitive function account implies that dorsal and ventral cellular recruitment differs depending on relevant behavioural demands, while the granularity account suggests similar recruitment dynamics regardless of the nature of the task performed. Here, we quantified cellular recruitment with the immediate early gene (IEG) *Arc* across the entire longitudinal CA1 axis in female and male rats performing spatial- and fear-guided memory tasks. Our results show that recruitment is greater in dorsal than ventral CA1 regardless of task or sex. This *experimentum crucis* leads to the strong inference that the granularity hypothesis for functional differences across the longitudinal axis in the rodent hippocampus is correct.

## Introduction

The hippocampus functionally differs across its longitudinal axis; exactly how to characterize this difference remains unclear. On one view, the dorsal and ventral aspect of the hippocampus support distinct behavioural processes [1-3]. Specifically, the dorsal region is said to be necessary for spatial navigation and memory, and the ventral region is important for emotional processing [2, 4]. An alternative view, largely informed by the observation that place field size increases from dorsal to ventral regions, posits instead that the hippocampal longitudinal axis varies in its granularity of representation, wherein spatial-temporal resolution of representation decreases along the axis, ventrally [5-7].

One approach to address differences in dorsal-ventral hippocampal functional properties is to examine cellular recruitment dynamics using electrophysiological, imaging, or genetic assays, such as immediate early gene (IEG) expression in tasks with different cognitive-behavioral demands. According to the behavioural function view, cellular recruitment will vary according to changes in task demands, such that dorsal regions will be most recruited during spatial navigation, and ventral regions in tasks that require emotional processing. By contrast, the granularity perspective predicts a descending gradient of cellular recruitment along the hippocampal longitudinal axis, regardless of the specific task being performed due to the lower number of cells needed to form a coarse representation of task state-space.

Work on cellular activation during open field navigation and dry-land memory tasks has demonstrated recruitment probability is greater in dorsal than ventral hippocampal subfields CA1-3 and the dentate gyrus [8, 9]. Recently, we found the same effect on activation in CA1 with the IEG *Arc* during memory-guided navigation in the Morris Water Task (MWT) [10]. One group also reported a similar gradient of expression in CA1 at the protein-level with the protein cFos [11]. Several groups have also reported that contextual fear conditioning augments *Arc* expression in dorsal and ventral hippocampal subfields and is necessary for contextual fear memory [12-15]. However, it remains unclear whether cellular recruitment probability across the hippocampal long axis depends on task or is invariant to changes in cognitive-behavioural demands.

To answer this question, we quantified cellular recruitment with the IEG *Arc* across the longitudinal axis of CA1 in female and male rats performing the MWT or context fear discrimination (CFD), which are both impaired following hippocampal damage [16-21]. We focused our behavioral measures on the MWT and CFD due to: 1) their popularity as behavioral assays of long-term memory; 2) differences in extra-hippocampal circuit mechanisms that support the expression of spatial and fear memory in the MWT and CFD, respectively [22-24]; 3) to extend our previous results on *Arc* mRNA expression in the MWT across sexes [10]. While these tasks may share emotional components during early performance, others have shown that stress responses in the MWT are elevated in early sessions but significantly reduced during later training [25]. Our group and others have also shown that regions associated with emotional processing such as the basolateral amygdala are not required for the expression of spatial memory in the MWT [26], but are required for CFD [27]. We further measured *Arc* mRNA expression as an indicator of population activity due to its established role in LTP, memory retrieval, and relatively precise temporal pattern of expression compared to other IEG transcripts and protein-level expression [28]. *Arc* has been used in numerous studies of cellular recruitment across the hippocampal long-axis and thus allows for comparison between our results and previous findings [8-10, 29]. Using this method, we designed a crucial experiment to adjudicate between two predicted outcomes based on the cognitive function and granular views of dorsal-ventral hippocampal activity, respectively: 1) *Arc* expression will be greater in dorsal than ventral CA1 in the MWT due to greater requirements of spatial processing during memory retrieval and, by contrast, greater in ventral than dorsal CA1 during CFD due to greater emotional processing in fear memory retrieval; 2) *Arc* expression will be greater in dorsal than ventral CA1 regardless of the task performed due to differences in the granularity of task representation across the longitudinal axis.

## Materials and Methods

### Subjects

The University of Lethbridge Animal Welfare Committee approved all procedures used in the present experiments, which also meet the Canadian Council of Animal Care guidelines. A total of 24 Long Evans rats weighing between 300 and 350 grams were used in the present experiments, including 12 females and 12 males (Charles River, Raleigh, NC). Previous work in our lab and others have determined such group sizes provide adequate statistical power for within-animal and across-group comparisons for behavioural and cellular levels of analysis [8, 10, 31, 32]. Following their arrival at the University of Lethbridge, animals were allowed at least 1 week to acclimatize to colony room conditions and were handled 5 minutes each day by the experimenter for 5 days before the start of the experiment.

### Experimental Design

To match behavioural experience across animals, male and female rats were equally divided into cohorts and trained in the MWT and CFD in counterbalanced order before final testing and perfusion (Figure 1). Our rationale for training animals both tasks was two-fold: 1) to ensure similar cumulative experience prior to final testing and perfusion that might otherwise affect plasticity and subsequent *Arc* transcription dynamics; 2) to ensure similar performance of groups in MWT and CFD performance. During training in the MWT, rats were transported from the animal colony to the behavioural testing room in covered cages on a cart. Surrounding the swimming pool were several posters on the walls, a table, and computer rack that served as allocentric cues to help animals navigate. During each session, animals were monitored, and relevant behavioural variables (latency and quadrant dwelling) were calculated using EthoVision XT software (Noldus) from an overhead behavioural camera. The MWT apparatus consisted of a 2-meter diameter pool filled with room temperature water (∼25 C) that was made opaque using white non-toxic tempura paint. On days 1-3, rats were given 8 trials (maximum 60 seconds) starting randomly from one of the four cardinal positions at the edge of the pool to locate a hidden platform approximately 5 cm beneath the water surface located in the center of the Northwest quadrant. If the animal did not locate the hidden platform within 60 seconds they were placed onto the platform by the experimenter. Animals were then allowed 10 seconds to remain on the platform before placement back into their holding cage by the experimenter for an approximate 5-minute intertrial interval. Following completion of the 8 swim trials each day, rats were returned to their colony room for approximately 24 hours prior to subsequent behavioural training or testing. During the final testing day in the MWT, animals were individually transported in covered cages and given 4 swim trials with a 2-minute intertrial interval for a total 10-minute testing period.

**Figure 1.**
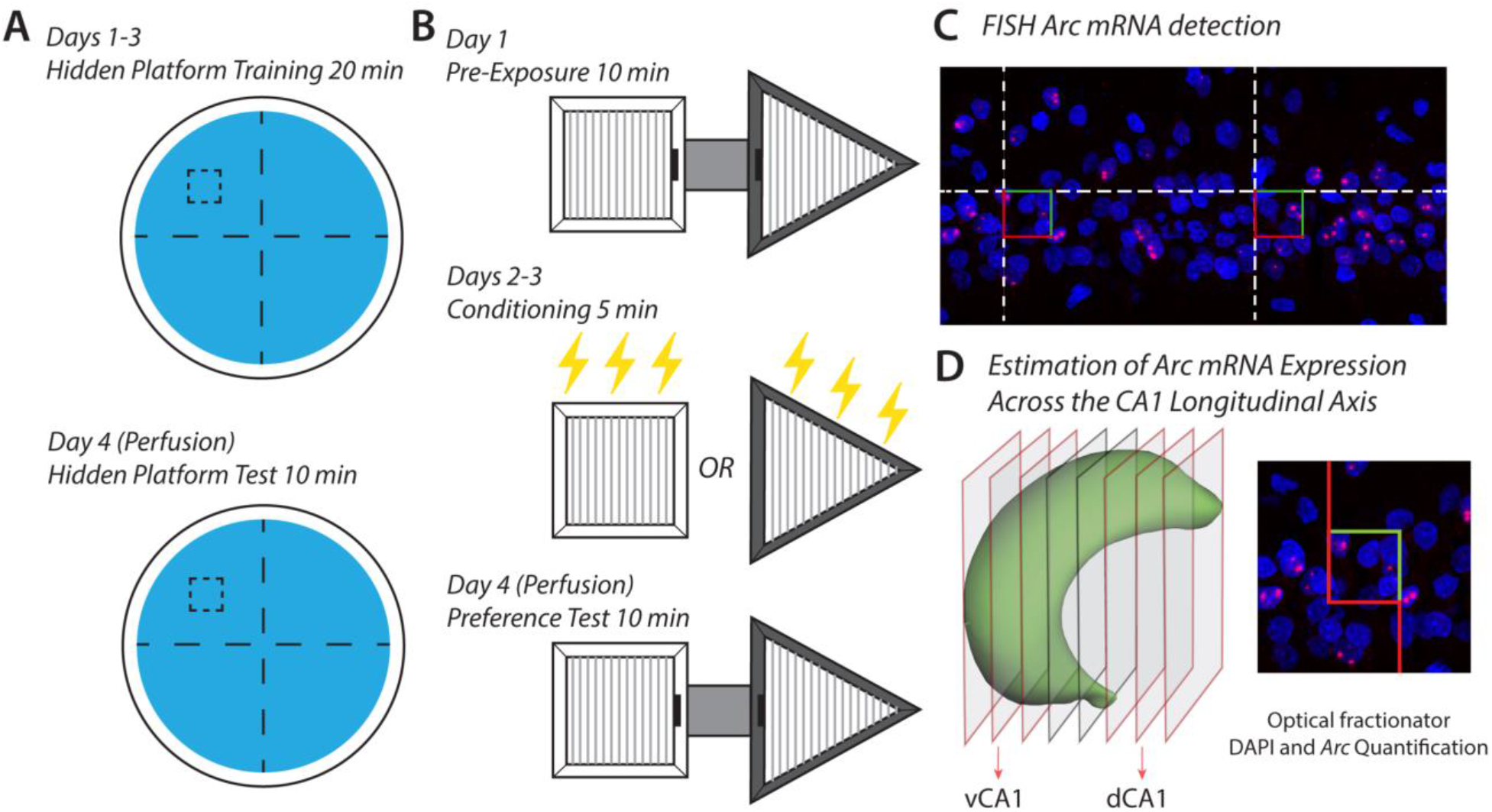
Experimental design to measure *Arc* mRNA expression across the CA1 long axis during MWT and CFD performance. A) Female and male rats were trained in the MWT using a 4-day procedure, wherein animals learned to swim to locate a hidden platform under the pool surface on days 1-3 and were given a 4-trial test on day 4. Half of the female and male groups were perfused approximately 5-10 minutes following testing in the MWT. B) Each animal was also trained in CFD, which began with 10 minutes of pre-exposure to both contexts connected by an alleyway on day 1. Following pre-exposure, animals were conditioned in counterbalanced order to shock-paired and unpaired contexts on days 2-3. Finally, animals were given a 10-minute test to examine preference for the unpaired context and avoidance of shock-paired context on day 4. C) Following testing in the MWT or CFD, animals were perfused, and brains were collected for FISH tissue processing. Brains were then sectioned on a freezing-sliding microtome at 40 μm thickness in a 12-section series, slide mounted, and stained with 4’,6’-diamidino-2-phenylindole (DAPI) and digoxygenin-conjugated antisense riboprobes for visualization of nuclear *Arc* mRNA expression (see Methods). A schematic stereological counting grid is overlaid to illustrate the approximate sampling frequency from CA1. D) Following tissue processing, DAPI and *Arc* expression were quantified using the optical fractionator method at 60X magnification on a confocal microscope according to the principles of systematic-random sampling. Estimates from dorsal and ventral CA1 subregions were then divided following quantification across the entire CA1 axis, wherein dorsal CA1 included sections anterior to -3.8 mm AP from bregma, and ventral sections included sections posterior to -5.2 mm AP from bregma [30].

In the CFD task, rats were individually transported in a covered holding cage by the experimenter to a room with several posters on the walls, a storage shelf, and the CFD apparatus, which consisted of two conditioning chambers (contexts) and connecting alleyway. One context was a black triangle that was 61 cm long, 61 cm wide, and 30 cm high with stainless steel rod flooring, and was scented with banana (Isoamyl Acetate, Sigma) located in a perforated pill bottle at the top right corner of the chamber. The other context was a white square that was 41 cm long, 41 cm wide, and 30 cm high with stainless steel rod flowing and was scented with Eucalyptus (Vic’s VapoRub©) located in a perforated pill bottle inserted through the top left corner of the chamber. During each session, animal behaviour was recorded from a tripod-mounted camera from underneath the apparatus through a transparent table. During pre-exposure on day 1 of the CFD task, animals were introduced to the apparatus through the connecting alleyway and allowed to freely explore both contexts for 10 minutes. On subsequent days 2 and 3, animals were conditioned in shock-paired and unpaired contexts in counterbalanced order. During unpaired conditioning, animals were placed in their unpaired context for 5 minutes and allowed to explore freely. For shock-paired conditioning, animals were transported to a distinct and separate room containing the same apparatus and placed into their paired context. The stainless-steel rod flooring was connected to a Lafayette Instrument Stimtek SGCG1 through a custom shock harness, and 2-second, 1.0 mA scrambled foot shocks were delivered at the 2^nd^, 3^rd^, and 4^th^ minute. After an additional 58 seconds, animals were removed from the shock-paired context and returned to their home cage. On day 4 of the CFD task, animals were returned to the original training room, and introduced to the apparatus through the connecting alleyway and allowed 10 minutes to explore both contexts. Dwell time in each context was calculated by a trained observer from video data during the pre-exposure and preference testing epochs and was defined as the presence of both forepaws in a context.

Following final testing in either the MWT or CFD, animals were returned to their holding cage for 1 minute, and then given an overdose intraperitoneal injection of sodium pentobarbital. They were then perfused 5-10 minutes after the completion of behavioural testing in either task, and then decapitated and had their brains extracted for subsequent tissue processing. This timeline was chosen based on previous studies demonstrating that behaviour-driven *Arc* expression is maximal 5-10 minutes after a learning or remembering episode [33].

### FISH tissue processing

The methods used for *Arc* visualization with FISH are identical to those in our previous report on *Arc* mRNA expression in the MWT [10]. Following fixation and sectioning at 50 µm thickness in a 12-section series using a freezing-sliding microtome, samples were stored at -80 C until FISH processing. *Arc* riboprobes were designed to detect intronic mRNA sequences, and thus nuclear rather than cytoplasmic expression. Primers flanking *Arc* intron 1, exon 2, and intron 2 were designed using online software (National Center for Biotechnology Information Primer-Blast; credit to A. M. Demchuk, University of Lethbridge). The exact sequences of the primers and base pair designations follow those of the GenBank accession number NC_005106: 5’-CTTAGAGTTGGGGGAGGGCAGCAG-3’ (forward primer, base pairs 2,022 - 2,045) and 5’-ATTAACCCTCACTAAAGGGCCCTGGGGCCTGTCAGA-TAGCC-3’ (reverse primer tagged with T3 polymerase binding site on 5’ end, base pairs 2,445-2,466). The polymerase chain reaction (PCR) was performed on a genomic rat DNA template using a Taq PCR Kit (New England Biolabs), and the subsequent PCR product was purified using a Qiagen PCR Purification Kit (Life Technologies, Inc.). The MAXIscript T3 transcription kit (Life Technologies, Inc.) and DIG RNA Labeling Mix (Roche Diagnostics) were used to generate DIG-labelled *Arc* intron-specific antisense riboprobes from PCR templates. Riboprobes were then purified with mini QuickSpin columns (Roche Diagnostics), and FISH was performed on slide-mounted tissue as previously described [10, 34, 35]. Briefly, DIG-labelled *Arc* riboprobe signal was amplified with anti-DIG-POD (1:300; Roche Diagnostics), Tyramide Signal Amplification (TSA) Biotin Tyramide Reagent Pack (1:100; PerkinElmer), and Streptavidin-Texas Red (1:200; PerkinElmer). Cell nuclei were then counterstained with DAPI (1:2,000; Sigma-Aldrich).

### Optical fractionator confocal stereology

The approach used for quantification of *Arc* and DAPI labels across the CA1 axis was identical to the methods described in Lee et al. (2019). Briefly, observers blind to experimental conditions of each animal quantified DAPI and *Arc* expression using the optical fractionator method in StereoInvestigator software (version 10.54, MBF Bioscience, VT) from confocal z-stack images collected on an Olympus FV1000 microscope equipped with Fluoview software (version 4.0, Olympus, Shinjuku, Japan). Unilateral traces of CA1 were created at 20X magnification on each section, and counting frames were automatically positioned according systematic-random sampling procedures with a 150 × 150 μm grid over CA1 traces. A series of seven z-stack images at 512 × 512-pixel resolution were collected at each sampling site with a 60X oil-immersion objective starting at the top of the section every 2 μm for a total 14 μm sampling distance in the z-plane. Image thresholds were set at 700 HV ± 20 and 550 HV± 20, respectively in DAPI and Texas Red channels, and kept constant across imaging each section series such that small *Arc* foci (2-3 pixels in diameter) and DAPI labels could be clearly identified. Digital z-stack images were then imported into StereoInvestigator software, such that the top image from each stack fell above and the final image below a 10-μm height of the optical dissector volume. *Arc* and DAPI were then counted according to optical fractionator inclusion–exclusion criteria at each cell’s widest point in a 30 × 30 × 10 μm fractionator probe [36].

### Statistical Analysis

Data were analyzed and visualized using the Prism by GraphPad statistical package, in addition to Numpy, Scipy, and Matplotlib libraries with Python 3.7 in Google Colab. A two-way or three-way ANOVA was used to examine main effects and interactions in behavioural and imaging data and *post hoc* Student’s t-test with Šidák correction following a significant interaction term. The variables considered as independent factors in these analyses included sex, quadrant (MWT), context (CFD), epoch, and CA1 subregion (dorsal or ventral). To perform Bayesian analysis, surrogate datasets were generated according to the granular and cognitive hypotheses as described in Results. This analysis and accompanying code can be found in the following Google Colab document: https://colab.research.google.com/. Following generation of surrogate data of dorsal and ventral recruitment probability, we calculated the gradient of recruitment as the difference between dorsal and ventral values in simulated data and from actual subjects. Normal distributions were then fit to each gradient in the MWT and CFD, and we calculated the BF as the ratio of the integrated product between the actual data and granular distributions versus cognitive hypothesis in each task. Using this method, we interpreted a BF from 10 to 30 as strong evidence for the granular hypothesis, and a BF score from 1/10 to 1/30 as strong evidence for the cognitive view.

## Results

Our behavioural results in the MWT and CFD show clear, successful learning in spatial and fear tasks for both male and female rats under the present training and test protocols. A two-way analysis of variance (ANOVA) of latency to locate the hidden platform in the MWT showed significant effects of day (*F*(3,66)=45.01, *p*<0.0001) and sex (*F*(3,66)=5.129, *p*=0.0337), but not a significant day x sex interaction (*F*(3,66)=0.08432, *p*=0.9684; Figure 2A), with female rats slower than males to reach the hidden platform. To ensure that the effect of sex on MWT performance was not an artifact of slower swim speed, we also examined the percentage of dwell time in target versus non-target pool quadrants across days (Figure 2B). A three-way ANOVA revealed day (*F*(3,66)=24.45, *p*<0.0001), quadrant (*F*(1,22)=464.9, *p*<0.0001), and sex (*F*(1,22)=11.50, *p* = 0.0026) as significant factors, in addition to significant sex x quadrant (*F*(1,22)=11.50, *p*=0.0026) and day x quadrant interactions (*F*(3,66)=24.45, *p*<0.0001). Thus, like some previous reports we found that males are somewhat faster and more accurate in allocentrically-based MWT navigation than females [37-40]. Despite effects of sex, post-hoc comparisons between target and non-target quadrants on day 4 showed significant preference for the target quadrant in both male (*t*=17.04, *p*<0.0001, DF=176) and female rats (*t*=14.01, *p*<0.0001, DF=176), demonstrating robust learning of the allocentric MWT in both males and females. Although we did find sex differences in the MWT, both cohorts performed similarly in CFD. A two-way ANOVA on total dwell time in paired and unpaired contexts during pre-exposure and preference testing revealed a significant effect of context (*F*(1,22)=4.927, *p*=0.0371), epoch (*F*(1,22)=12.63, *p*=0.0018), and context x epoch interaction (*F*(1,22)=14.31, *p*=0.0010), but no effect of sex (*F*(1,22)=2.639, *p*=0.1185), context x sex (*F*(1,22)=0.9049, *p*=0.3518), epoch x sex (*F*(1,22)=0.1768, *p=0*.*6782*), or context x epoch x sex interaction (*F*(1,22)=1.059, *p*=0.3145). To control for possible differences in time spent dwelling in the connecting alleyway during the preference test, we also examined the percent dwell time in paired and unpaired contexts; excluding time spent in the connecting alleyway. A three-way ANOVA on percent dwell time in CFD showed significant effects of context (*F*(1,22)=7.205, *p*=0.0135) and context x epoch interaction (*F*(1,22)=24.47, *p*<0.0001), but no effect of epoch (*F*(1,22)=0.9995, *p*=0.3283), sex (*F*(1,22)=1.000, *p*=0.3281), sex x context (*F*(1,22)=0.4861, *p*=0.4930), sex x epoch (*F*(1,22)=1.000, *p*=0.3281), or sex x context x epoch interaction (*F*(1,22)=0.6604, *p*=0.4251). *Post hoc* comparisons also showed significant preference for the unpaired context during preference testing in both males (*t*=3.824, *p*=0.0015, DF=88) and females (*t*=5.854, *p*<0.0001, DF=88). These results suggest that both male and female cohorts expressed robust spatial- and fear-guided behaviour to promote *Arc* expression.

**Figure 2.**
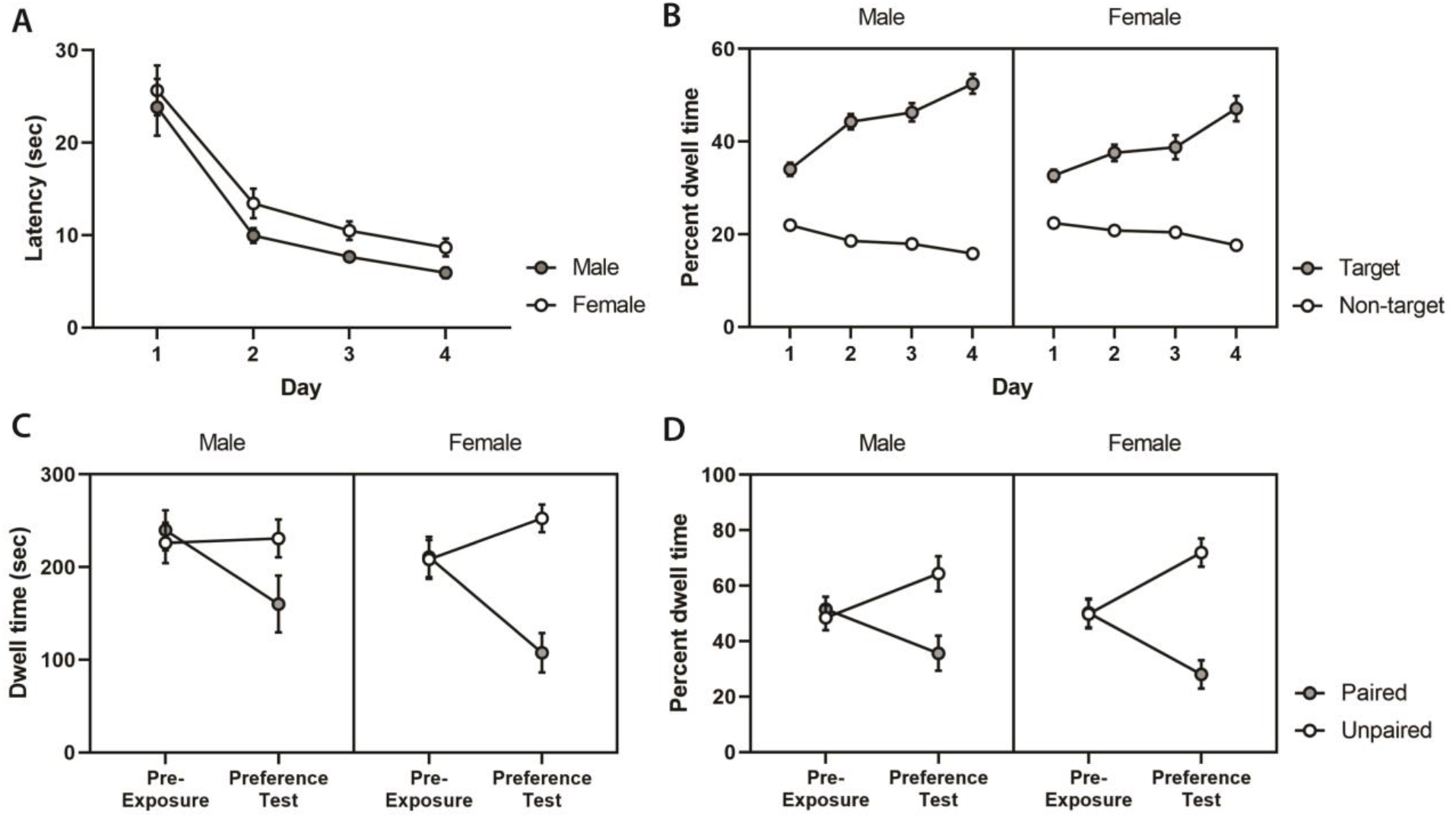
Female and male behavioural performance in the MWT (A, B) and CFD (C, D). A) Average latency to reach the hidden platform decreased across training (1-3) and test (4) days for both females and males, with males somewhat faster to reach the hidden platform. B) Percent dwell time in target and non-target quadrants showed a strong increase in preference for the target quadrant across training and testing, suggesting that animals effectively learned to navigate to the hidden platform in the MWT. C) Total dwell time in shock-paired (paired) and unpaired contexts in the CFD task shows similar preference for both contexts during pre-exposure, and preference for the unpaired context after fear conditioning in female and male animals. D) Controlling for variation in time spent in the connecting alleyway during pre-exposure and preference testing, percent dwell time in paired and unpaired contexts was calculated excluding time spent in the connecting alleyway. Our results demonstrate a similar percent dwell time in the paired and unpaired contexts during pre-exposure, and a strong preference for the unpaired context following conditioning, with no clear effect of sex on CFD performance.

Following behavioural testing in the MWT and CFD, we quantified *Arc* expression using design-based confocal stereology across the entire CA1 longitudinal axis. DAPI-stained cell bodies and *Arc* transcription foci were quantified using the optical fractionator method from confocal z-stacks according to the principles of systematic-random sampling (see Methods for details). We estimated DAPI and *Arc* populations across CA1, including dorsal (anterior to -3.8 mm AP from bregma; Figure 3A) and ventral (posterior to -5.2 mm from bregma; Figure 3A) subregions, and transformed these estimates into a single *Arc*:DAPI metric of cellular recruitment probability. A three-way ANOVA of estimated DAPI and *Arc* populations across CA1 (Figure 3B) showed significant effects of label (*F*(1,20)=1281, *p*<0.0001) and sex (*F*(1,20)=7.023, *p*=0.0154), but no effect of task (*F*(1,20)=0.08018, *p*=0.7800), label x sex (*F*(1,20)=2.203, *p*=0.1533), task x sex (*F*(1,20)=0.1407, *p*=0.7115), task x label (*F*(1,20)=0.5850, *p*=0.4533), or task x sex x label interaction (*F*(1,20)=0.7562, *p*=0.3948). Notably, our DAPI estimates closely resemble previous reports of cell number across CA1 (mean DAPI = 418313) using a design-based stereological approach [10, 41]. Following transformation of population estimates into *Arc*:DAPI recruitment probability, a two-way ANOVA showed no effect of task (*F*(1,20)=0.08314, *p*=0.3727), sex (*F*(1,20)=3.083, *p*=0.0944), or task x sex interaction (*F*(1,20)=0.7737, *p*=0.3895) on recruitment probability when the entire CA1 population was considered (mean male=0.1728; mean female=0.2021; Figure 3C). Notably, the magnitude of *Arc* expression observed in CA1 was comparable to levels reported in several studies of open field navigation and dry-land memory tasks, and well above levels observed in home-cage control animals, which is typically between .04 - .08 proportion of cells across studies in dorsal and ventral CA1 [9, 29, 32]. Such levels of home cage activation have been reliably reported across research groups with different quantification methods, and strongly support that the levels of cellular recruitment in the present study are behaviourally-driven [8, 9, 29, 32, 42]. To address whether cellular recruitment differs across axis, task, or sex, estimates were divided into dorsal and ventral CA1 *Arc*:DAPI recruitment probabilities (Figure 3D). A three-way ANOVA showed a significant effect of axis (*F*(1,20)=77.35, *p*<0.0001), but no effect of task (*F*(1,20)=0.2344, *p*=0.6335), sex (*F*(1,20)=1.772, *p*=0.1981), axis x task (*F*(1,20)=0.03415, *p*=0.8553), sex x task (*F*(1,20)=0.2616, *p*=0.6146), axis x sex (*F*(1,20)=0.3346, *p*=0.5694), or axis x task x sex interaction (*F*(1,20)=0.3325, *p*=0.5706). Our results thus do not support the cognitive hypothesis on recruitment probability across the CA1 longitudinal axis.

**Figure 3.**
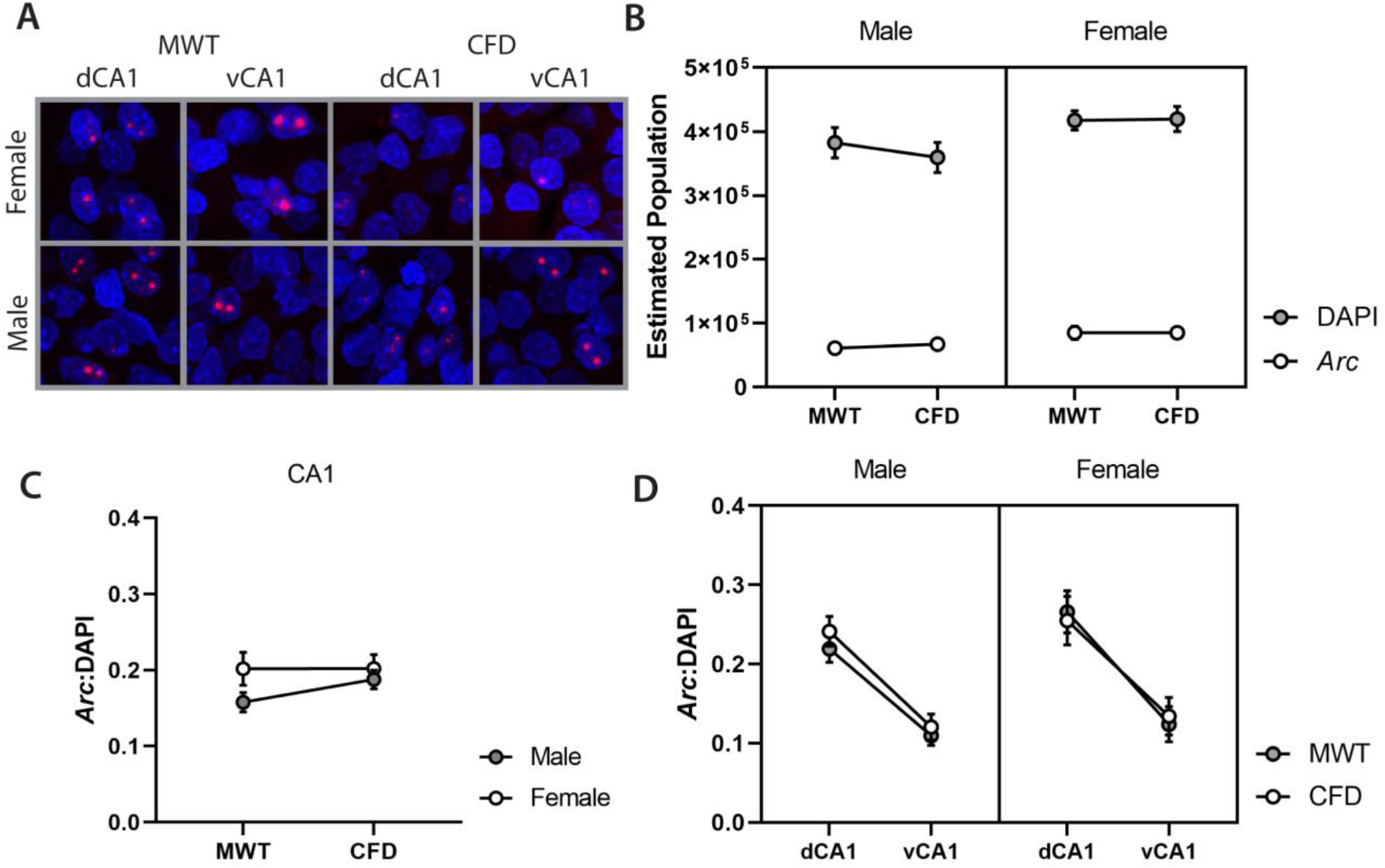
Quantification of DAPI and *Arc* across the CA1 longitudinal axis in male and female rats performing the MWT and CFD. A) Randomly chosen example images of DAPI and *Arc* labelling in dorsal and ventral CA1 in male and female animals that performed the MWT or CFD. B) Estimated populations of DAPI and *Arc* across CA1 in males and females were similar previous studies using design-based stereology in the rat hippocampus [10, 41]. C) Following estimation of DAPI and *Arc* populations, estimates were transformed into *Arc*:DAPI recruitment probabilities for the entire CA1 population. We did not find any differences between males and females in the MWT or CFD *Arc*:DAPI recruitment probabilities when examining the entire CA1 population. D) *Arc*:DAPI recruitment probabilities were divided into dorsal and ventral CA1 estimates for males and females performing the water task to examine evidence for the granular and cognitive hypotheses. These results show a clear effect of axis in males and females performing both the MWT and CFD, wherein dorsal CA1 has greater recruitment probability than ventral CA1 regardless of task or sex differences.

The cognitive and granular accounts of cellular recruitment make distinct predictions about the gradient of *Arc* expression across the CA1 long axis in the MWT and CFD. Namely, the cognitive model predicts opposite gradients of cellular recruitment in the two tasks, and the granular model predicts the same gradient across tasks. Importantly, we anticipate opposite outcomes in the CFD task according to each hypothesis, but the same outcome in the MWT. While the above analysis rejects the cognitive account of long axis cellular recruitment, we aimed to directly compare evidence for each view using a Bayesian approach [43]. To this end, we generated surrogate data resembling CA1 estimates of cellular recruitment in each task according the granular and cognitive hypotheses, wherein the granular model is described by a decaying beta probability distribution ventrally between α = [5.00, 1.00] and β = [10.00, 12.00], and the cognitive model is a task-dependent beta probability distribution from α = [5.00, 1.00] and β = [10.00, 12.00] in the MWT, and the opposite gradient between α = [1.00, 5.00] and β = [12.00, 10.00] in CFD. In addition, we approximated cellular recruitment of home cage controls in our model based on previous reports from several labs that found a similar level of recruitment across dorsal and ventral CA1 [9, 29, 32, 42], with model parameters set to α = [1.00] and β = [15.00] for both dorsal and ventral CA1, which yielded an average recruitment of approximately 0.05. We then computed the average dorsal and ventral recruitment probabilities from each model and determined the task-dependent recruitment gradient as the difference between dorsal and ventral recruitment in our *Arc*:DAPI dataset and surrogate data (Figure 4A). Finally, we fit a gaussian likelihood distribution to each dorsal-ventral gradient and computed the Bayes Factor (BF) as the ratio of the integrated product of the actual data from both experimental groups versus the distribution of surrogate home cage data to compute a task-related BF. We then calculated BF for hippocampal long-axis models in both tasks from the integrated product of the actual and granular likelihood distributions, versus the actual data and the cognitive likelihood distributions (Figure 4C). This provides a single BF that describes the ratio of fit between the actual data and each hypothesized distribution, wherein a BF between 10 and 30 was considered as strong evidence for and the granular hypothesis, and a BF between 1/10 and 1/30 as strong evidence for the cognitive view. In comparison to home cage surrogate data (Figure 4B), BF above 10 or below 1/10 is considered strong evidence for a task-related cellular recruitment gradient. Using this approach, we found strong evidence for a task-related gradient (BF = 13.68), suggesting that the pattern of CA1 recruitment we observed is not explained from results expected in home cage control animals. Comparing models of hippocampal long-axis function, we found similar evidence for both hypotheses in the MWT (BF = 1.07; Figure 4C), and strong evidence for the granular hypothesis in CFD where the predictions of each account differ (BF = 23.09; Figure 4C). The combined results of these analyses thus favor the granular account of cellular recruitment across the CA1 long axis over the cognitive function view.

**Figure 4.**
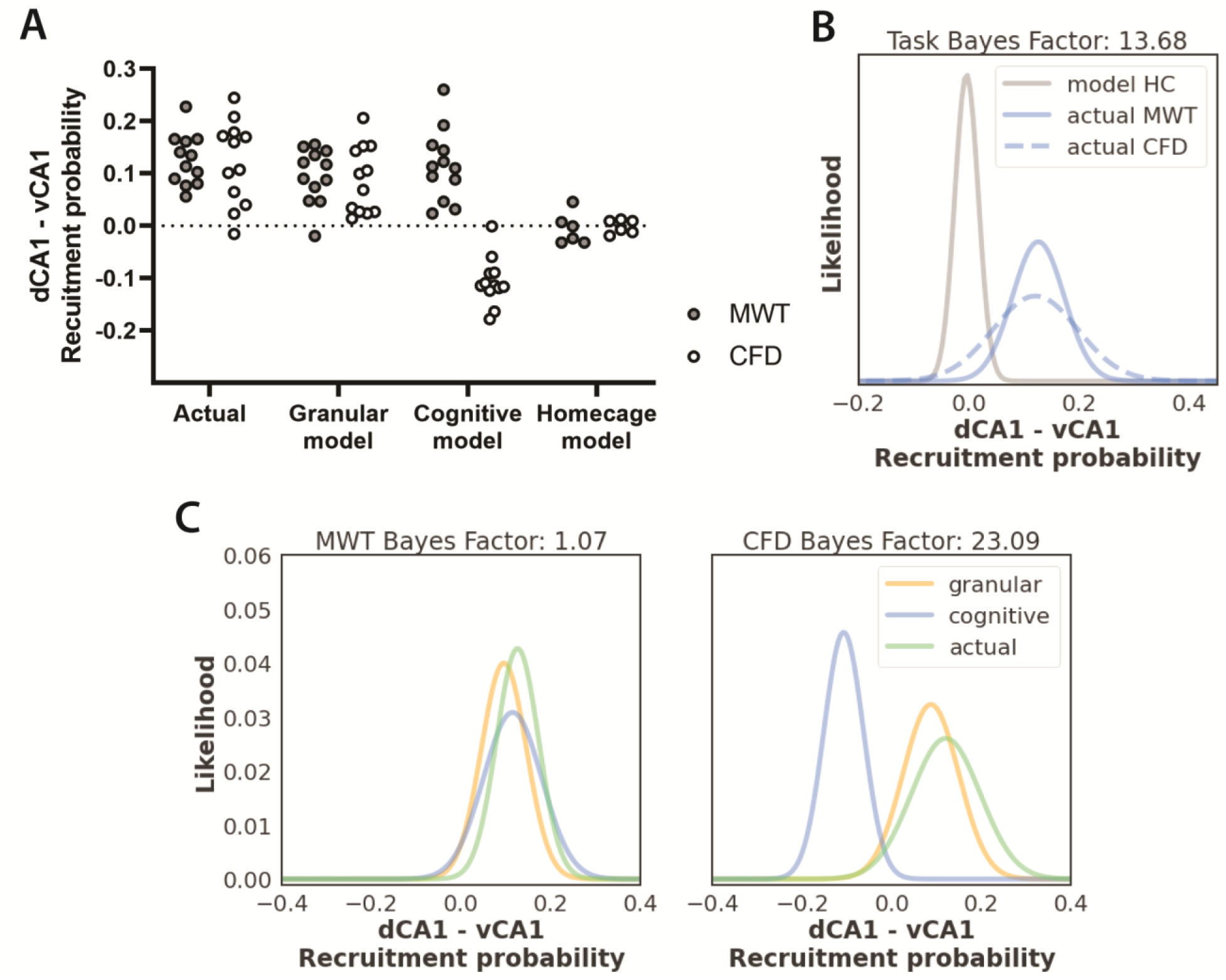
Bayesian analysis of differential CA1 recruitment probability across the hippocampal long axis in the MWT and CFD. A) Female and male data were combined from the MWT and CFD and the gradient of recruitment probability was calculated as the difference between dorsal and ventral CA1 *Arc*:DAPI estimates. Surrogate data were generated according to home cage control observations in previous studies [8, 9, 29, 32], and the granular and cognitive hypotheses of dorsal and ventral CA1 cellular recruitment. We then computed the difference between dorsal and ventral estimates from surrogate and actual data in the MWT and CFD to generate model-based recruitment probability gradients. B) Normal distributions were fit to actual and surrogate home cage data to compute a task-related BF as the ratio of the integrated product of actual distributions compared to the home cage control distribution, which demonstrated the gradient of cellular recruitment we observed is behaviourally driven. C) Normal distributions were also fit to surrogate and actual dataset from the MWT and CFD, and the BF was computed as the ratio between the integrated products of the actual and surrogate distributions according to the granular versus the cognitive hypothesis. Using this approach, we observed greater evidence for the granular over the cognitive view of the hippocampal long axis function.

## Discussion

In the present experiments, we have found that cellular recruitment probability is greater in dorsal than ventral CA1 during performance of the MWT and CFD in both male and female rats. While previous work has demonstrated the same effect in using the IEG *Arc* during open-field navigation and memory retrieval [8, 9], our results are the first to demonstrate such a gradient of activity across the hippocampal long-axis in a fear-based task. Further, the present findings replicate our previous result that cellular recruitment is greater in dorsal than ventral CA1 during navigation in the MWT [10], and bear great similarity to the pattern of recruitment in tasks that have no clear aversive component [8, 9, 29]. These findings across several studies and research groups suggest that the dorsal-ventral gradient of activity in CA1 may be an invariant property of cellular recruitment in this region. Our results are also the first to our knowledge that demonstrate such recruitment dynamics are shared across sexes. Despite differences in performance between males and females in the MWT, we did not find differences in their pattern cellular recruitment. Finally, our analysis comparing these results against the granular and cognitive function hypotheses on cellular recruitment demonstrate clear support for the granular view on hippocampal long-axis function (Figure 4). Future work should examine how such gradients might also differ across additional hippocampal subregions, including the dentate gyrus (DG), CA3, and the subiculum. If the granularity hypothesis is correct for all subregions, we anticipate the same gradient of cellular recruitment in the MWT and CFD, with inter-regional differences in overall levels of activation, with the DG, CA3, CA1, and subiculum showing increasing levels of recruitment, respectively, similar to previous open-field navigation studies [8, 9, 44].

Evidence from lesion, electrophysiological, functional imaging, and IEG studies supports that the hippocampus functionally differs across its longitudinal axis [1, 4, 5, 8, 17]. While arguments have been made from these observations in support of either a cognitive or granular account of this difference, the present results on cellular recruitment in CA1 lend greater support to the granular hypothesis. Several observations from navigation, memory, and cognitive mapping literature inform this view. Of these findings, perhaps the most characterized is the gradient in place field size that increases across the hippocampal long axis ventrally. Several groups have found that while average place field diameter is often <1 meter in dorsal hippocampal regions, field diameter in the ventral pole can range up to 10 meters [5, 45]. In keeping with this observation, recent comparisons of dorsal and ventral hippocampal lesions in the MWT have shown that while dorsal lesions impair precise localization of goal locations, ventral damage impairs coarse localization in the MWT [16-18]. In relation to cellular recruitment, it is anticipated that cells in the dorsal hippocampus are more likely to be recruited due to a greater number of small place fields that would be required to tesselate a space than those with large fields in ventral hippocampus [6, 7]. Indeed, multiple groups have found that place cells in the dorsal hippocampus have multiple fields in large environments [46-50]. Recently, Harland et al. [47] also reported that place cells recorded in the dorsal hippocampus during navigation in a two-dimensional “megaspace” (>18 m^2^) express multiple fields of varying sizes that tend to increase with scale of the environment. They also found a negative correlation between field number and size, suggesting that perhaps the granularity of representation might be related to the recruitment probability. However, the relationship between place field size and recruitment probability across the hippocampal long axis is not well understood. Future experiments to evaluate the granularity view might consider manipulations of room size or spatial sampling during open field navigation as in Witharana et al. [44] and measuring cellular recruitment across the hippocampal long-axis. Despite the present support from these experiments for the granular view of the hippocampal longitudinal axis, it is important to consider evidence that might also favour the cognitive view.

The cognitive function view stems from considering anatomical projections and functional cell types that differ in dorsal and ventral regions. While dorsal CA1 and subiculum have strong efferent projection to the retrosplenial, rhinal, and post-rhinal cortices, ventral CA1 and subiculum strongly project to the rhinal and medial prefrontal cortices (mPFC), basal and central amygdala, ventral striatum, and lateral hypothalamus [1, 4, 51]. As a result, information represented across the hippocampal long axis projects to distinct targets and receives input from many of these regions. Previous reports have shown that synchronized activity between the ventral hippocampus and mPFC modulates anxiety behaviour [52], and recent calcium imaging experiments have also revealed functional cell types that respond to shock and anxiety in ventral hippocampus that project monosynaptically to the basal amygdala and lateral hypothalamus, respectively [51, 53]. However, despite the presence of anxiety and shock-responsive cells in ventral hippocampus, recent studies also point to the existence of shock-responsive cells in dorsal CA1 [54, 55]. Further, several groups have found that many cells in dorsal CA1 are responsive to reward [48, 56-59], and optogenetic activation of such cells results in reward-seeking behaviour [57]. While some studies have reported differential impairments of dorsal and ventral hippocampal lesions in tasks that require emotional regulation, such as contextual fear conditioning, some also demonstrate that dorsal and ventral lesions similarly impair performance in such tasks, including non-spatial delay tone conditioning [2, 60-62]. Indeed, our group and others have also shown that CFD is impaired following either dorsal, ventral, and complete hippocampal damage [16, 19, 63, 64]. While some evidence supports that behavioural variables are differently coded across the hippocampal long axis, the causal relationship between dorsal and ventral regions with spatial and emotional behaviour may not be mutually exclusive as the cognitive hypothesis suggests.

Another possible explanation of differences that we have observed in *Arc* expression across the CA1 longitudinal axis could be related to cellular excitability and intrinsic recruitment propensity. In dorsal CA1, several studies have reported that cellular recruitment does not follow a Poisson process (random draw with replacement), but instead is gamma or log-normally distributed [44, 48, 49]. Recently, Lee et al. (2020) found that most cells in CA1 are virtually “silent” and have no place fields across multiple, large environments, while some cells have single fields, and a minority of cells have multiple fields in multiple environments – in keeping with recruitment statistics of hippocampal cells in multi-room IEG studies [44]. Importantly, Lee et al. [48] also found that cellular excitability is positively related to the propensity of cells to express fields for multiple places, environments, rewards, and across time. Cellular excitability and resultant propensity likely have a direct role in cellular recruitment probability and coding sparsity. While previous whole-cell patch recordings have revealed that ventral CA1 neurons are more intrinsically excitable than in dorsal CA1 [65, 66], it remains unclear whether the excitability-propensity relationship is constant across the hippocampal long axis, and whether propensity-based recruitment distributions are similar or differ. Based on the present results and previous studies of cellular recruitment [8, 10, 44], we anticipate fewer overall cells to have fields in ventral than dorsal CA1 possibly due to differences in afferent projection and less dendritic length and surface area in ventral CA1 [65]. However, those cells with the propensity to have fields would be more excitable in ventral than dorsal CA1. This would suggest that the gamma-distributed process of recruitment differs across the hippocampal long axis, although this has not been examined experimentally. Future work might therefore explore the relationship between excitability, propensity, and cellular recruitment across the hippocampal longitudinal axis, and its potential role in determining the precision of hippocampal representation and encoding of behavioural variables.

## Acknowledgements

We thank Valérie Lapointe, Aubrey Demchuk, Nhung Hong, and Maurice Needham for their assistance with this project.

